# Heterozygous variants in the mechanosensitive ion channel *TMEM63A* result in transient hypomyelination during infancy

**DOI:** 10.1101/682179

**Authors:** Huifang Yan, Guy Helman, Swetha E. Murthy, Haoran Ji, Joanna Crawford, Thomas Kubisiak, Stephen J. Bent, Jiangxi Xiao, Ryan J. Taft, Adam Coombs, Ye Wu, Ana Pop, Dongxiao Li, Linda S. de Vries, Yuwu Jiang, Gajja S. Salomons, Marjo S. van der Knaap, Ardem Patapoutian, Cas Simons, Margit Burmeister, Jingmin Wang, Nicole I. Wolf

## Abstract

Mechanically activated (MA) ion channels convert physical forces into electrical signals. Despite the importance of this function, the involvement of mechanosensitive ion channels in human disease is poorly understood. Here we report heterozygous missense mutations in the gene encoding the MA ion channel TMEM63A that result in an infantile disorder resembling a hypomyelinating leukodystrophy. Four unrelated individuals presented with congenital nystagmus, motor delay, and deficient myelination on serial scans in infancy, prompting the diagnosis of Pelizaeus-Merzbacher (like) disease. Genomic sequencing revealed all four individuals carry heterozygous missense variants in the pore-forming domain of TMEM63A. These variants were confirmed to have arisen *de novo* in three of the four individuals. While the physiological role of TMEM63A is incompletely understood, it is highly expressed in oligodendrocytes and it has recently been shown to be a mechanically activated (MA) ion channel. Using patch clamp electrophysiology, we demonstrated that each of the modelled variants results in strongly attenuated stretch-activated currents when expressed in naïve cells. Unexpectedly, the clinical evolution of all four individuals has been surprisingly favorable, with substantial improvements in neurological signs and developmental progression. In the three individuals with follow-up scans after four years of age, the myelin deficit had almost completely resolved. Our results suggest a previously unappreciated role for mechanosensitive ion channels in myelin development.

---

Leukodystrophies are genetic disorders primarily affecting brain white matter characterized by the abnormalities they demonstrate on magnetic resonance imaging (MRI) [1–6]. MRI pattern recognition often helps to obtain an MRI-based diagnosis in leukodystrophies and enables targeted genetic testing [7, 8], but genetic heterogeneity may require a more comprehensive approach. Next generation sequencing (NGS) has rapidly improved the diagnosis of previously elusive or novel phenotypes with whole exome sequencing (WES) and whole genome sequencing (WGS) now directly responsible for implicating a large number of genes with leukodystrophies, greatly reducing the number of unsolved cases [2, 9–12].

Myelination is a protracted developmental process that progresses in infancy following a fixed pattern and is almost complete by the age of 24 months. It can be followed using serial MRI scans [4, 13]. A severe deficit of myelin observed on time-separated scans is classified as a hypomyelinating leukodystrophy, a large and genetically heterogeneous group among the leukodystrophies [14]. Clinical presentation is often in early infancy with nystagmus and axial hypotonia, evolving to ataxia and spasticity, with motor development more affected than cognitive function. Alternatively, presentation later in life with a milder clinical picture is also possible [15,16]. The prototype of hypomyelinating leukodystrophies is Pelizaeus-Merzbacher disease (PMD, MIM: 312080) due to alterations of *PLP1* (OMIM:300401), encoding the structural myelin protein, proteolipid protein 1 [17]. Under the age of 24 months hypomyelination cannot be diagnosed on MRI with a single exam [14]. However, in the context of profound myelin deficit and a compatible clinical presentation, hypomyelinating leukodystrophy may be suspected in the first year of life and appropriate investigations can be started. Without a definitive diagnosis, MRI should be repeated to distinguish hypomyelination from delayed myelination.

Here we report on four individuals who presented with a severe myelin deficit observed on serial MRIs, indistinguishable from PMD in the infantile stage (Figure 1), which unexpectedly resolved on follow-up scans, with parallel clinical improvement. Families 1 & 2 were investigated as part of an on-going study on the Amsterdam Database of Leukoencephalopathies to unravel the genetic cause of unclassified leukodystrophies. Families 3 & 4 were identified in Peking University First Hospital (Beijing, China). Institutional ethics approval was granted at both participating centres and written informed consent secured from the guardians of these individuals. WGS and WES were performed on family trios (proband, biological mother, and biological father), for Families 1 & 2 and Families 3 & 4, respectively.

**Figure 1.**
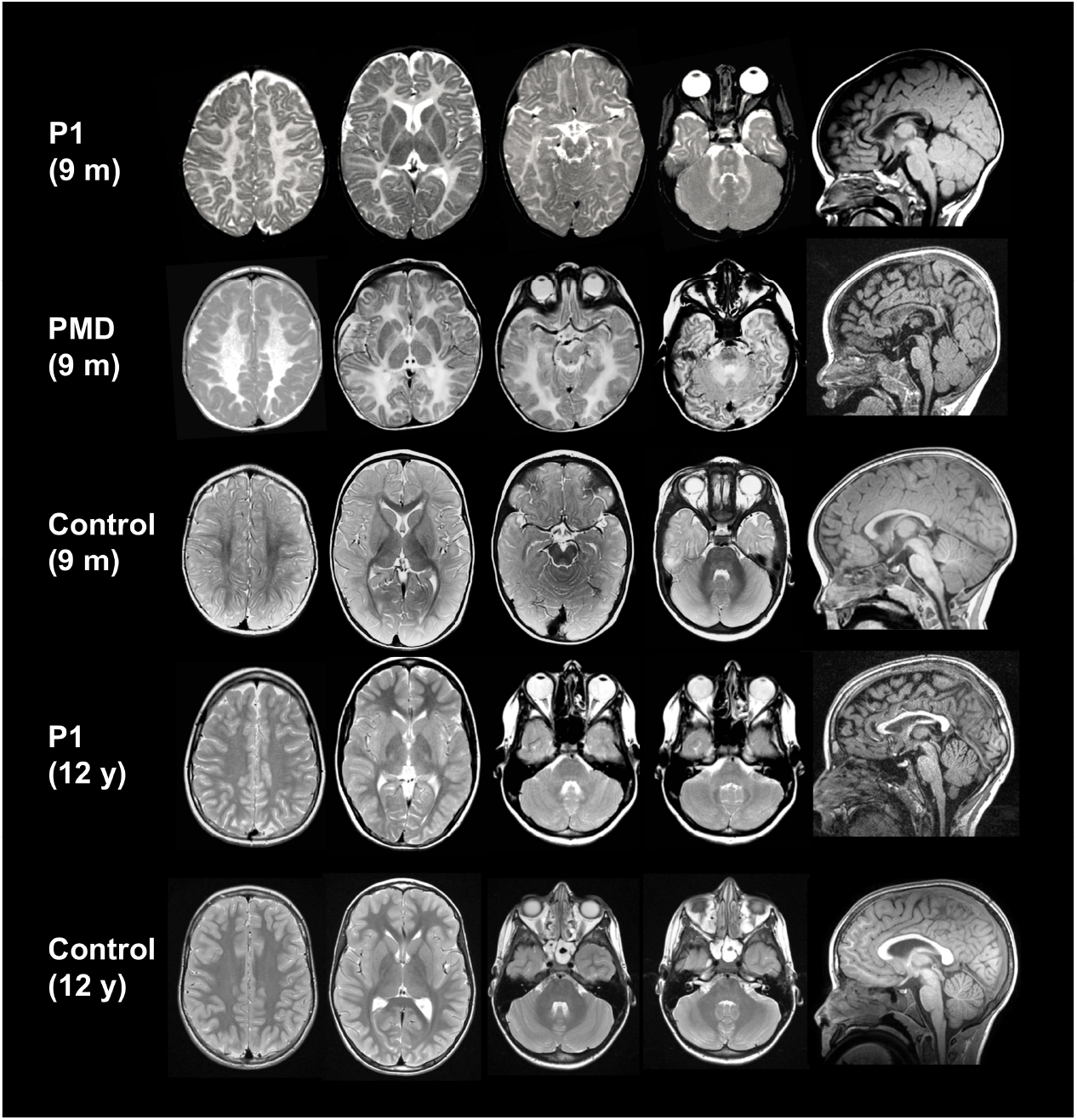
MRI evolution of individual 1. Axial T_2_-weighted images and a mid-sagittal T_1_-weighted image of individual 1 at 9 months and 12 years of age, shown in comparison with age-matched controls and a individual with PMD aged 9 months. At 9 months, MRI findings are indistinguishable from PMD, with supra- and infratentorial white matter signal that is diffusely elevated on the T_2_-weighted images. The corpus callosum is thinner than normal. At age 12 years, supratentorial white matter signal has normalized, but there is still mild hyperintensity of the cerebellar white matter and the middle cerebellar peduncles.

Individual 1 is a 21-year-old male of European descent who was born at term without complications after an uneventful pregnancy. At age 2 weeks, parents noted a pendular nystagmus, increasing with fixation and when agitated. At the age of 2 months, he was an alert baby who could follow objects, but had a tendency towards opisthotonus. In the following months, his development slowly progressed, but at age 8 months, he was not able to sit because of poor balance and had head titubation. He could not grasp objects well due to intention tremor and dysmetria. He could move around by rolling over. His nystagmus had persisted and was now fast. Brain MRI at age 9.5 months showed severe myelin deficit, with no normal myelin signal of both supra- and infratentorial white matter at T_1_- and T_2_-weighted images (Figures 1 and 2), similar to an age-matched child with PMD due to *PLP1* duplication (Figure 1). At 1 year of age, he was able to sit relatively stable and started to pull himself up. Language development was normal.

**Figure 2.**
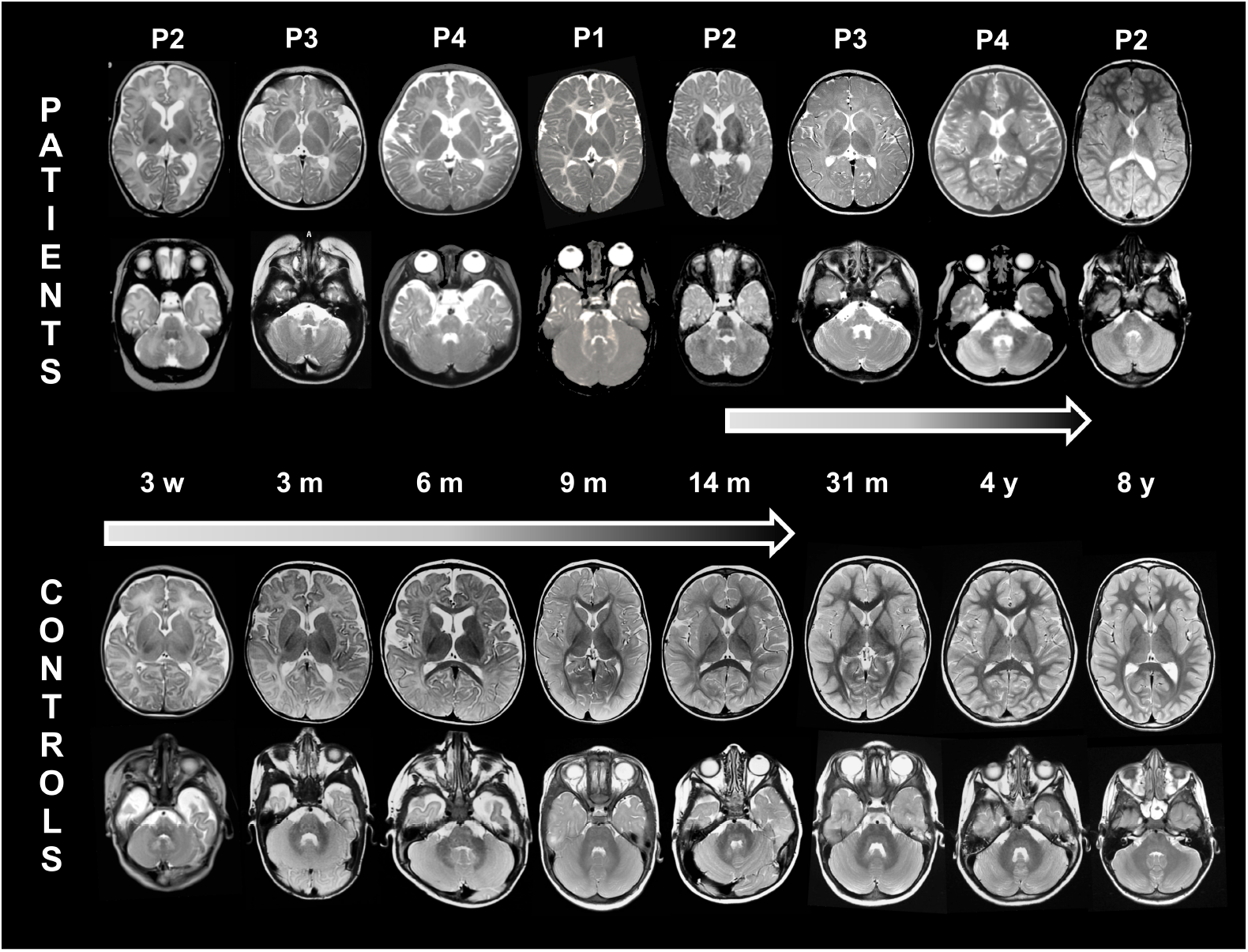
Transient hypomyelination in individuals with *TMEM63A* variants. Serial MRIs (axial T_2_-weighted images) of the 4 individuals, from age 3 weeks (left) to age 4 years (right), alongside corresponding images of age-matched healthy children. The normal images of individual 2 at age 8 years and the MRI of individual 1 at age 12 years (see also Figure 1 for this individual) are not depicted here. Myelination shows small improvements only at age 31 months and is completed by age 4 years, whereas it is continually progressing in healthy children, to be virtually completed at age 2 years.

He walked without support at age 20 months. At 5 years, neurological examination showed only mild ataxia; his nystagmus had almost disappeared. He had low vision and mild myopia. Fundoscopy was normal. At age 7 years, he had an episode of vertigo, without nystagmus. Both brain CT and CSF testing were normal, and vertigo improved within 2 to 3 weeks. At 12 years of age, brain MRI showed normal myelin signal with the exception of the middle cerebellar peduncles and the cerebellar white matter, which were mildly T_2_ hyperintense (Figure 1). At age 13 years, he had very mild ataxia with suboptimal tandem gait and standing on one leg. By age 16 years, he had developed optic atrophy. At that age, he developed Crohn’s disease, which is stable with treatment. Without cognitive problems, he visited regular school and is now a student at university.

Individual 2 is a 17-year-old male of European descent born at 38 weeks gestational age after a pregnancy complicated by diabetes mellitus treated with insulin. Delivery was by vacuum extraction, and birth weight was 4700g. Immediately post-partum, he had to be resuscitated (Apgar scores 1/1/1), but did not need intubation. On day 2 of life, he developed epileptic seizures, necessitating treatment and artificial ventilation. Brain ultrasound was repeatedly normal. After one week, he was seizure-free and discharged at age two weeks. Brain MRI at age 3 weeks showed no evidence of hypoxia related brain lesions, but the normal myelin signal in the posterior limb of the internal capsule (PLIC) was absent (Figure 2). He received antiepileptic treatment during 3 months. Although abnormal brain-stem auditory evoked potentials (BAEP) were observed, his parents thought his hearing was fine. He was first seen in clinic at 4 months, where a pendular nystagmus was noted, accompanied by low axial muscle tone and poor head control with titubation. At age 14 months, brain MRI showed diffuse T_2_-hyperintense signal of both supra- and infratentorial white matter (Figure 2). His development slowly progressed, and he was able to walk without support at age 17 months. Language development was slightly delayed. At age 7 years, he had mild gait and appendicular ataxia. His nystagmus had resolved. He went to regular school, but mild learning difficulties became increasingly evident. His IQ was 87 at age 7 years. At 8 years, MRI revealed resolution of hypomyelination. In addition, he had persistent ductus arteriosus, which had to be closed at age 18 months. Hypospadia was corrected at age 8 months. He lacked several teeth and had mild myopia. Family history was significant for both parents and several siblings. His mother has diabetes mellitus and his father died at age 60 years from a brain tumor. The father’s vision was good, and it is not known whether he had nystagmus or delayed development in early life. Five older siblings of the affected individual in this family all declined genetic testing. Two brothers have mild cognitive problems, one sister has diabetes mellitus type I, and the other two sisters are healthy. Two of the siblings, now 2 and 5 years of age, had transient nystagmus, which was not further investigated.

Individual 3 is a 5-year-old female and is the second child of an unrelated Chinese couple and has a healthy older sister. Pregnancy and delivery were uneventful. Her birth weight was 3050g. Nystagmus was observed 10 days after birth and had resolved by the end of the first year of life. Motor developmental delay was first noted at age 7 months when she still could not hold her head. Gesell development scale (Chinese version) assessed at 7 months showed mild development delay. She started to walk without support at age 26 months. Currently, she is able to run and jump, but falls easily. Compared to her motor abilities, cognitive development was relatively spared. She started to smile socially at age 2 months and recognized relatives by 4 months. Language expression and understanding were normal, but her pronunciation was suboptimal. With continuous improvement and no regression, she attended regular preschool with normal performance. Hearing and vision were clinically intact. Physical examination performed at 7 months of age demonstrated only mild hypotonia. Brain MRI performed at 3 and 31 months of age implied diffuse myelin deficit in both the supra- and infratentorial brain white matter (Figure 2), compatible with a hypomyelinating leukodystrophy.

Individual 4, now 4 years old, is the second child of unrelated Chinese parents. He was born uneventfully at term by elective cesarean section. His birth weight was 3600g. Physiological jaundice was observed and spontaneously subsided after one month. Nystagmus was noted after birth and resolved by 14 months of age. At the age of 6 months, vision was decreased and myopia was diagnosed. Fundoscopy was normal. Visual evoked potentials (VEP) performed at age 6 and 49 months were delayed. His motor development was delayed, with some catching up after the first year of life. Head control was achieved at age 10 months, sitting without support at age 16 months, standing on his own at age 24 months and walking without support at age 36 months. He started to speak at age 39 months and currently is able to speak long sentences, but with unclear pronunciation. Gesell development scale (Chinese version) assessed at age 6 months, 17 months and 50 months showed variable impairment in all domains of development. On follow-up he had made developmental progression. His hearing was clinically normal, though BAEP performed at age 6 months and 49 months had demonstrated delayed conduction of binaural listening pathways in the brainstem and increased hearing thresholds. Myopia was still present. Brain MRI performed at age 6 months and 13 months showed diffuse myelin deficit in the cerebral white matter, which had resolved at age 50 months (Figure 2).

In all of these individuals, results of routine investigations and metabolic testing were normal. PMD was suspected clinically, but *PLP1* dosage was normal as were targeted NGS of 115 leukodystrophy-related genes (including *GJC2*) or individual sequencing of *POLR3A*/*POLR3B* in Individual 2. Complete case descriptions are available in the Supplemental Data.

The earliest MRI in our cohort was performed at age 3 weeks in Individual 2. The earliest MRIs for all affected individuals, in contrast to healthy newborns, showed no myelin signal in the posterior limb of the internal capsule (PLIC) (Figure 2). There was no progression of myelination in the ensuing months on T_2_-weighted images with diffuse T_2_-hyperintense signal of both supra- and infratentorial white matter, including T_2_-hyperintense white matter in the cerebellum and the pyramidal tracts and medial lemniscus in the brain stem (Figure 2), in contrast to controls where these structures are myelinated as early as 3 months (Figure 2). Myelination had progressed slightly at 14 months (Individual 2, Figure 2) when the PLIC, the genu of the corpus callosum, and the frontal periventricular white matter developed a signal isointense with the cortex on T_2_-weighted images and the cerebellar white matter became almost isointense with the cerebellar cortex. Myelination had normalized in all individuals tested, the earliest by 4 years (Individual 4, Figure 2). The last MRI available at 12 years (Individual 1) showed mildly T_2_-hyperintense signal of the cerebellar white matter and the cerebellar peduncles, reflecting (permanent) white matter involvement (Figure 1).

In two independent studies, NGS detected heterozygous missense variants in *TMEM63A* (NM 014698.2) in each of the four families, including one recurrent variant. The candidate gene was posted on GeneMatcher [18]. *De novo* variants were identified in Individual 1 [c.1699G>A p.(Gly567Ser)], Individual 3 [c.1385T>A;p.(Ile462Asn)], and Individual 4 [c.503G>A p.(Gly168Glu)]. Individual 2 shared the same variant as Individual 1 [c.1699G>A p.(Gly567Ser)], but in this case it was found to be paternally inherited. The father of Individual 2 is deceased and records from his childhood were unavailable to determine possible clinical similarities in the infantile period. Individual 2 was also found to have compound heterozygous missense variants in *SLC6A9*, encoding a glycine transporter, that were initially investigated as candidates. However, glycine uptake assays indicated these variants did not have an obvious impact on transporter activity (results not shown), excluding these variants as candidate pathogenic alleles. Individual 4 also has a single *de novo* variant [c.1936G>A p.(Ala646Thr)] in *GRIA1* (NM 001258022.1) [19], which has been reported with development delay without MRI abnormalities. Interestingly, milestone achievement was more severely delayed in Individual 4 (Table 1). Therefore, we suspect that variants in *TMEM63A* and *GRIA1* may both contribute to his phenotype. No other candidate variants emerged from the genomic analysis of these individuals.

**Table 1.** Clinical Characteristics

| Individual | 1 | 2 | 3 | 4 |
| --- | --- | --- | --- | --- |
| **Variant** | Gly567Ser | Gly567Ser | Ile462Asn | Gly168Glu |
| **Gender** | male | male | female | male |
| **Current Age** | 21Y | 17Y | 5Y | 4Y |
| **Nystagmus** |  |  |  |  |
| Age at onset | 14D | 1D | 10D | 1D |
| Age resolved | 5Y | 7Y | 12M | 14M |
| **Development** |  |  |  |  |
| Walking without support | 20M | 17M | 26M | 36M |
| Language development | Normal | Slightly delayed | Normal | Delayed |
| **MRI** |  |  |  |  |
| Myelin deficit (age) | 9.5M | 21D,14M | 3M, 31M | 5M, 13M |
| Myelin Normalization (age) | 12Y* | 8Y* | NA | 4Y* |
| **Myopia** | + | + | - | + |
| **Findings at last neurological examination** | Optic atrophy,<br>mild ataxia | Mild ataxia | Unclear<br>pronunciation | Bilateral<br>babinski sign,<br>unclear<br>pronunciation |
| **Other** | Crohn's disease,<br>one self-limiting<br>episode of vertigo | Patent ductus<br>arteriosus, hypospadia,<br>abnormal BAEP<br>and VEP | - | Abnormal<br>BAEP and VEP |

Note: Y, years; D, days; M, months; NA, not available; \*, Earliest exam available; BAEP, brain-stem auditory evoked potential; VEP, visual evoked potential.

The *TMEM63A* variants identified in this study are not present in the 1000 Genomes or gnomAD human population databases and are all predicted to be damaging by multiple prediction tools: SIFT, Polyphen-2, MutationTaster, M-Cap, Condel and PROVEAN (Supplemental Table). The three mutated residues are also highly conserved across vertebrate species (Figure 3).

**Figure 3.**
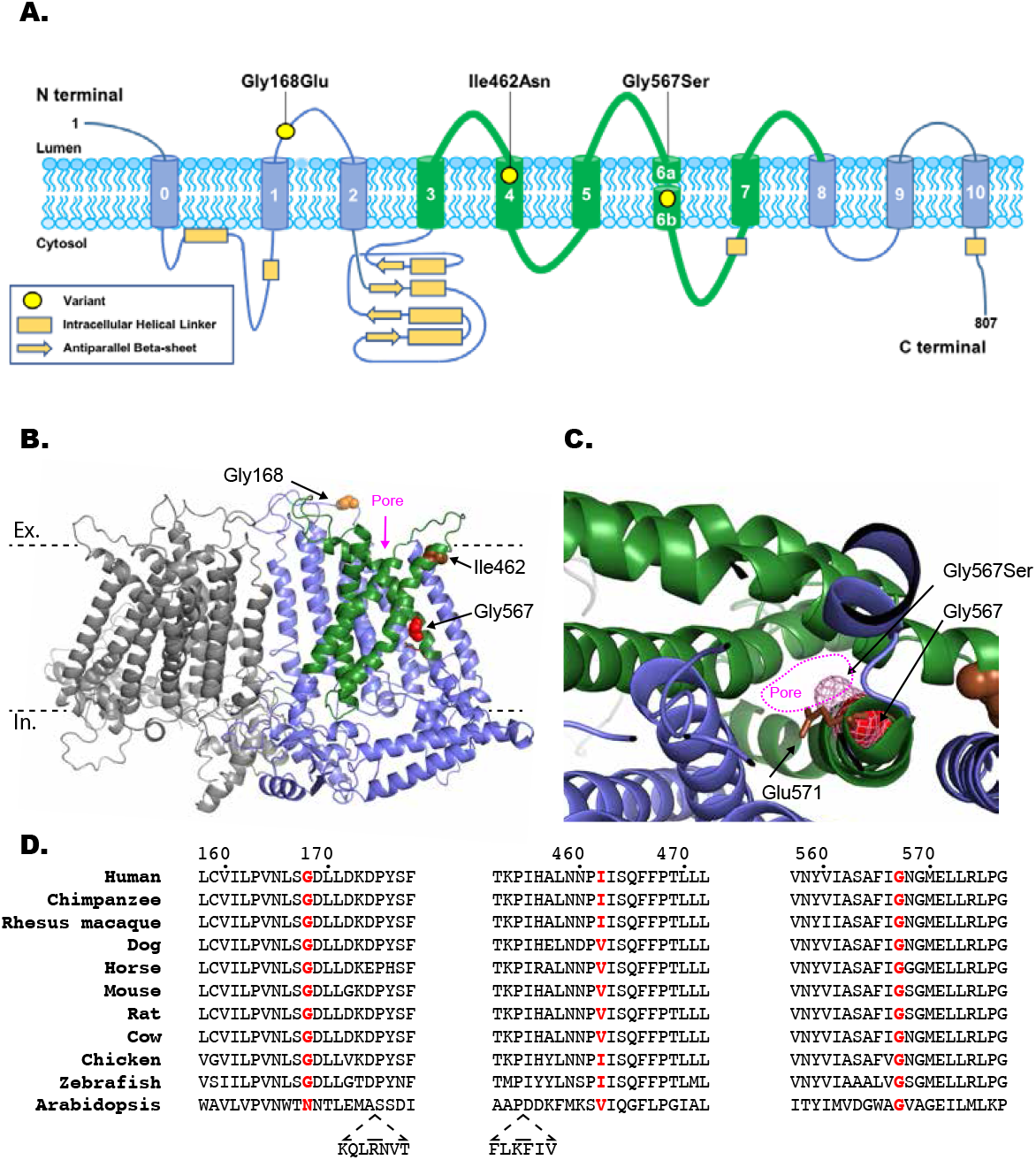
Protein structure and variant conservation. A) Location of the three identified variants shown on a schematic representation of TMEM63A protein. B) and C) Predicted structure TMEM63A protein based on cryo-EM structure of Arabidopsis homologue OSCA1.2 (PDB: 6MGV). Model generated using SWISS-MODEL [20, 21]. Residues altered by the newly found variant are shown as spheres. The approximate location of the extracellular (Ex) and intracellular (In) surfaces of plasma membrane indicated by dashed black lines. The five pore lining transmembrane helices are shown in green [20]. C) Section of TMEM63A structural model looking through pore region towards the extracellular domain. Gly576 residue shown as red spheres, pink mesh indicated the predicted position the serine side-chain would occupy in the p.Gly576Ser mutant protein. Conserved residue Glu571 (shown as brown sticks) that has been shown to reduce stretch activated ion conductance when mutated to alanine in OSCA1.2 can be seen adjacent to Gly576 [20]. D) Clustal alignment of TMEM63A protein from 10 vertebrate species and the Arabidopsis homologue OSCA1.2. The residues altered in the presented individuals are highlighted in red.

*TMEM63A* is highly expressed in oligodendrocytes. Mouse *Tmem63a* shows significantly higher expression in myelinating oligodendrocytes and in microglia (http://web.stanford.edu/group/barres_lab). In a large-scale mouse phenotyping study, a small but significant proportion of *Tmem63a* null mice demonstrate gait abnormalities (www.mousephenotype.org/data/experiments?geneAccession=MGI:2384789).

Using a cryo-EM structure of the TMEM63A homologue OSCA1.2 from Arabidopsis, we generated a model of TMEM63A protein structure using SWISS-MODEL (Figure 3b) [20, 21]. The Gly168 residue sits in an intracellular loop between helix 1 and helix 2 while both Ile462 and Gly567 are located within pore lining transmembrane helices of TMEM63A. Interestingly Gly567 resides on the internal face of the pore one helix turn from the conserved residue Glu571 that has been shown to reduce stretch activated ion conductance when mutated to alanine in OSCA1.2 [20].

We introduced the variants seen in these individuals by site directed mutagenesis into TMEM63A clones purchased from Origene (NM_014698, Cat No.: RC206992). Wild-type and mutant constructs were sub-cloned into an IRES_mCherry vector and transfected into PIEZO1-knockout Human Embryonic Kidney 293T cells. Using a previously described cell-based stretch-activated channel activity assay [20, 22], we characterised the impact of each of the three variants on TMEM63A channel activity (Figure 4). Mechanical stimulation, induced by the application of negative pressure in the recording electrode, elicited stretch-activated currents in cells transfected with wild-type TMEM63A. However, cells transfected with each of the three *TMEM63A* variants (Gly168Glu: N=7; Ile462Asn N=9; Gly567Ser: N=7) failed to induce stretch-activated currents (Figure 4B), suggesting that these variants cause a loss of function.

**Figure 4.**
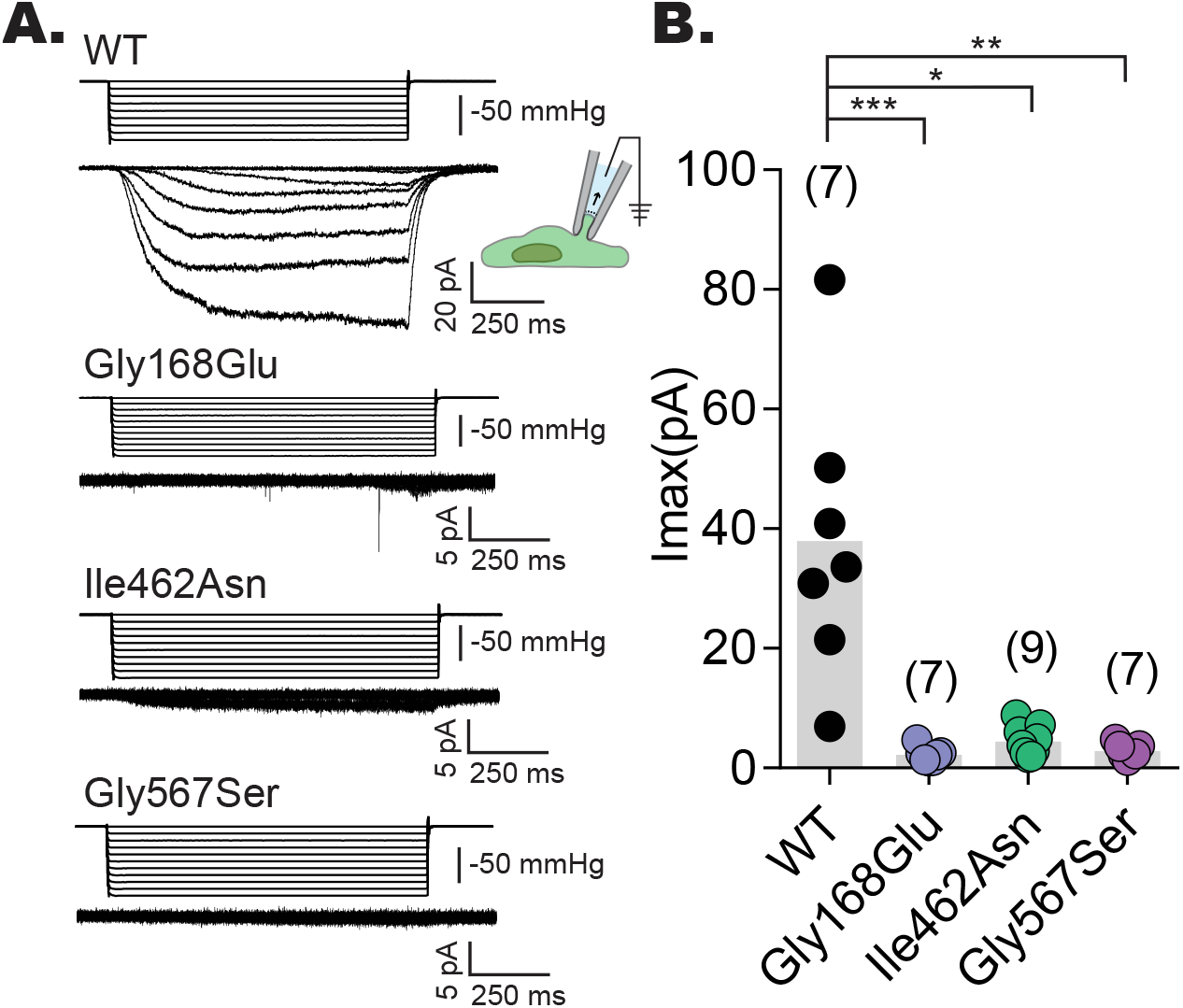
Mutant constructs fail to induce MA currents in HEK-P1KO cells. A) Representative stretch-activated currents induced by negative pipette pressure from cells transfected with wild-type or the indicated mutant construct. Corresponding pressure stimulus is illustrated above the current trace. Vertical scale bar: 50 mmHg. B) Maximal stretch-activated currents recorded from cells expressing wild-type TMEM63A (N = 7) or the indicated variant of TMEM63A Gly168Glu (N = 7), Ile462Asn (N = 9), and Gly567Ser (N = 7). ***p=0.0002, *p=0.0411, **p=0.0031, Dunn’s multiple comparison test.

The data presented here associate heterozygous variants in *TMEM63A* with a leukodystrophy with transient myelin deficiency during infancy. All four cases had clinical presentations suggestive of PMD or PMLD at an early age with nystagmus and delayed motor development and MRIs demonstrating a pattern suggestive of hypomyelination (present on serial MRIs in three individuals and profound myelin deficiency on single MRIs in the other infant). However, clinical evolution in all four individuals has been surprisingly favorable, with resolution of nystagmus, near-normalization of neurological signs and developmental progression. Concurrent with the improvement in neurological signs, MRI showed unexpected normalization. In two individuals the myelin deficit appears to have completely resolved and in one individual only mild abnormalities persist. The remaining individual, last imaged at 31 months, also displayed improvement of myelination.

The distinction between hypomyelination and delayed myelination can be challenging. PMD is a severe neurologic disorder characterized by hypomyelination with lasting cognitive and motor deficits. Individuals with PMD may achieve small developmental gains, but do not acquire sitting without support and demonstrate progressive spasticity and gradual deterioration typically by 10 – 12 years of age. Delayed myelination can be a non-specific finding across a spectrum of infantile onset neurologic disorders. Profoundly delayed myelination with slow improvement and even complete resolution of myelin deficiency on MRI, has been observed in Allan-Herndon-Dudley syndrome (AHDS), due to variants in *MCT8* encoding a thyroid hormone transporter [4, 23]. However, in AHDS, individuals typically remain severely affected, and clinical signs do not improve even when there are radiologic changes that suggest normalization of the myelin deficit [23, 24]. Temporally, neither MRI findings nor clinical development were compatible with PMD or AHDS in individuals with variants in *TMEM63A* despite their early presentation, thus we characterized this presentation as a transient myelin deficiency.

TMEM63A, also named CSC1-like protein 1, was initially suggested to be a hyperosmolarity activated ion channel [25, 26]. More recently it was demonstrated that TMEM63s are MA ion channels that elicit stretch-activated currents when expressed in naïve cells [22]. The results described here clearly demonstrate that each of the hypomyelination associated variants result in a loss of TMEM63A function either due to direct effect on channel gating or due to compromised protein folding and trafficking. This suggests a previously unappreciated role for *TMEM63A* in early myelin development. How disruption of this activity results in the apparently temporary myelin deficit observed in these individuals is unclear. *TMEM63A* has two highly similar homologues *TMEM63B* and *TMEM63C*. It is possible that developmental and tissue specific expression of these homologues provide some compensation for the loss of *TMEM63A* activity [22].

Finally, our findings add *TMEM63A* to a growing list of leukodystrophies caused by heterozygous variants, the majority of which arise *de novo*. Disorders that arise sporadically are refractory to techniques such as linkage analysis that permit an estimation of a molecular cause of disease. Trio NGS has demonstrated success in these cases, with heterozygous variants in *CSF1R*, *TUBB4A*, and *TMEM106B* found to cause a prominent leukodystrophy [9, 27, 28].

Further monitoring of these individuals described in this paper will be necessary to better understand the natural course of the disease related to heterozygous *TMEM63A* variants and to reveal whether there is a risk of slow regression over time. We propose a new phenotype of infantile-onset transient hypomyelination caused by heterozygous *TMEM63A* variants, characterized by clinical and radiological normalization in childhood, making this prognostically important information for affected families. It also emphasizes that myelination can evolve normally even when it is initiated later than usual, giving hope for therapeutic trials in hypomyelinating disorders.

## Supporting Information

A supplemental text file has been provided and includes a full description of methods and expanded clinical descriptions of the described individuals.

## Supplemental Table

**Characteristics of three variants in *TMEM63A***. Variant *in silico* prediction scores for all three variants found in this study.

## Acknowledgments

The authors would like to thank the affected individuals and their families. We thank Dr. Truus E.M. Abbink, Carola van Berkel and Nienke L. Postma for their invaluable assistance.

