## Supplemental Data for "Heterozygous variants in the mechanosensitive ion channel *TMEM63A* result in transient hypomyelination during infancy"

### Supplemental Materials & Methods

*Subjects*

Families 1 and 2 were investigated as part of an on-going study on the Amsterdam Database of Leukoencephalopathies to unravel the genetic cause of unclassified leukodystrophies. Families 3 and 4 were identified in Peking University First Hospital (Beijing, China). Approval from the institutional ethical committees of the participating centers was obtained and written informed consent secured from the guardians. Affected individuals were examined by neurologists at their primary care centers. Imaging data were reviewed by N.I.W. Genomic DNA specimens were obtained from circulating leukocytes using standard procedures.

*Genomic Analysis*

WGS and WES were performed on family trios (proband, biological mother, and biological father), for Families 1 & 2 and Families 3 & 4, respectively. For WGS, sequencing was performed using 2x150-nucleotide paired-end reads on an Illumina X10 by IIlumina Cambridge Ltd. Alignment of reads was performed using the Burrows-Wheeler Aligner.[28] For WES, a SureSelect Human All Exon V6 (Agilent, US) was used for exome capture. Sequencing was performed using 2x150-nucleotide paired-end reads on an Illumina X10 by Wuxi NextCODE Genomics (Shanghai) Co., Ltd. Alignment of reads was performed using the Sentieon software package. Variant calling for WGS and WES was performed using GATK HaplotypeCaller v3.7.[29] Variant annotation was performed using SnpEff v4.3m[30] and a custom script was prepared for variant filtration and prioritization. Candidate variants were posted on GeneMatcher.[31]

*Cell constructs and Electrophysiology.*

Laboratory studies were performed as previously described.[19; 21] In brief, TMEM63A clones were purchased from Origene (NM_014698, Cat No.: RC206992) and variants were introduced by site directed mutagenesis. WT and mutant constructs were sub-cloned into an IRES_mCherry vector and transfected into PIEZO1-knockout Human Embryonic Kidney 293T cells (HEK-P1KO, original HEK293T cell RRID: CVCL_0063)) using lipofectamine 2000 (Invitrogen).

Patch clamp recordings were performed between 16 to 48 hours after transfection using Axopatch 200B amplifier in a cell-attached configuration. External solution used to zero the membrane potential consisted of (in mM): 140 KCl, 1 MgCl2, 10 glucose and 10 HEPES (pH 7.3 with KOH). Recording pipettes were filled with a standard solution of 130 NaCl, 5 KCl, 10 Hepes, 1 CaCl2, 1 MgCl2, 10 TEA-Cl, pH 7.3. Membrane patches were stimulated with 1 second pulses of negative pressure, at a gradient of 10 mmHg, through the recording electrode using Clampex controlled pressure clamp HSPC-1 device (ALA-scientific). Electrophysiology data was analyzed using Clampfit 10.6 and data were plotted using GraphPad Prism6.

### Supplemental Note: Case Reports

*Individual 1*

This male individual, now 21 years old, was born at term without complications after an uneventful pregnancy. His parents, of European descent, are unrelated, and two younger brothers are healthy. At age 2 weeks, parents noted a pendular nystagmus, increasing with fixation and when agitated. At the age of 2 months, he was an alert baby who could follow objects, but had a tendency towards opisthotonus. In the following months, his development slowly progressed, but at age 8 months, he was not able to sit because of poor balance and had head titubation. He could not grasp objects well due to intention tremor and dysmetria. He could move around by rolling over. His nystagmus had persisted and was now fast. Brain MRI at age 9.5 months showed severe myelin deficit, with no normal myelin signal of both supra- and infratentorial white matter at $T_1$- and $T_2$-weighed images (Figures 1 and 2), similar to an age-matched child with PMD due to *PLP1* duplication (Figure 1) and clearly different from a healthy child. At 1 year of age, he was able to sit relatively stable and started to pull himself up. Language development was normal. He walked without support at age 20 months. At 5 years, neurological examination showed only mild ataxia; his nystagmus had almost disappeared. He had low vision and mild myopia. Fundoscopy was normal. At age 7 years, he had an episode of vertigo, without nystagmus. Both brain CT and CSF testing were normal, and vertigo improved within 2 to 3 weeks. At 12 years of age, brain MRI showed normal myelin signal with the exception of the middle cerebellar peduncles and the cerebellar white matter, which were mildly $T_2$ hyperintense (Figure 1). At age 13 years, he had very mild ataxia with suboptimal tandem gait and standing on one leg. By age 16 years, he had developed optic atrophy. At that age, he developed Crohn's disease, which is stable with treatment. Diagnostic evaluation included *PLP1* gene analysis and metabolic testing, with normal results. Without cognitive problems, he visited regular school and is now a student at university.

*Individual 2*

This male individual, now 17 years old, was born at 38 weeks gestational age after a pregnancy complicated by diabetes mellitus treated with insulin. Delivery was by vacuum extraction, and birth weight was 4700g. Immediately post partum, he had to be resuscitated (Apgar scores 1/1/1), but did not need intubation. On day 2 of life, he developed epileptic seizures, necessitating treatment and artificial ventilation. Brain ultrasound was repeatedly normal. After one week, he was seizure-free and discharged at age two weeks. Brain MRI at age 3 weeks showed no evidence of hypoxia related brain lesions, but the normal myelin signal in the posterior limb of the internal capsule was absent (Figure 2). He received antiepileptic treatment during 3 months. Although abnormal brain-stem auditory evoked potentials (BAEP) were observed, his parents thought his hearing was fine. He was first seen in clinic at 4 months, where a pendular nystagmus was noted, accompanied by low axial muscle tone and poor head control with titubation. This clinical presentation in combination with abnormal BAEP raised the suspicion of PMD, but genetic testing of *PLP1* did not confirm this diagnosis. At age 14 months, brain MRI showed diffuse $T_2$-hyperintense signal of both supra- and infratentorial white matter (Figure 2). His development slowly progressed, and he was able to walk without support at age 17 months. Language development was slightly delayed. At age 7 years, he had mild gait and appendicular ataxia. His nystagmus had resolved. He went to regular school, but mild learning difficulties became more and more evident. His IQ was 87 at age 7 years. At 8 years, MRI revealed resolution of hypomyelination. In addition, he had persistent ductus arteriosus, which had to be closed at age 18 months. Hypospadia was corrected at age 8 months. He lacked several teeth and had mild myopia. Sequencing of *POLR3A* and *POLR3B* was normal. Regarding family history, his parents were of European descent and not related. His mother has diabetes mellitus. His father died at age 60 years from a brain tumor. The father's vision was good, and it is not known whether he had nystagmus or delayed development in early life. The boy has five older siblings who declined further genetic testing. Two brothers have mild cognitive problems, one sister has diabetes mellitus type I, and the other two sisters are healthy. Two of the siblings children, now 2 and 5 years of age, had transient nystagmus, which was not further investigated.

*Individual 3*

This female individual, now 5 years old, is the second child of an unrelated Chinese couple and has a healthy older sister. Pregnancy and delivery were uneventful. Her birth weight was 3050 g. Nystagmus was observed 10 days after birth and had resolved by the end of the first year of life. Motor developmental delay was first noted at age 7 months when she still could not hold her head. Gesell development scale (Chinese version) assessed at 7 months showed mild development delay. She started to walk without support at age 26 months. Currently, she is able to run and jump, but falls easily. Compared to her motor abilities, cognitive development was relatively spared. She started to smile socially at age 2 months and recognized relatives by 4 months. Language expression and understanding was normal, but her pronunciation was suboptimal. With continuous improvement and no regression, she attended regular preschool and had normal performance. Hearing and vision were clinically intact. Physical examination performed at 7 months of age demonstrated no other abnormal neurological signs other than mild hypotonia. Brain MRI performed at 3 and 31 months of age implied diffuse myelin deficit in both the supra- and infratentorial brain white matter (Figure 2), compatible with a hypomyelinating leukodystrophy. Routine investigations were normal. Due to the combination of clinical and MRI findings, Pelizaeus-Merzbacher-like disease (PMDL, MIM:608804)) was suspected, but mutations in 115 leukodystrophy-related genes including *GJC2* were excluded by multiplex ligation-dependent probe amplification (MLPA) and targeted NGS (Kangso Medical Inspection, Beijing) and *PLP1* dosage was normal.

*Individual 4*

This boy, now 4 years old, is the second child of unrelated Chinese parents. He was born uneventfully at term by elective cesarean section. His birth weight was 3600g. Physiological jaundice was observed and spontaneously subsided after one month. Nystagmus was noted after birth and resolved by 14 months of age. At the age of 6 months, vision was decreased and myopia was diagnosed. Fundoscopy was normal. Visual evoked potentials (VEP) performed at age 6 and 49 months were delayed. Severe developmental delays in the following milestones were reported (age in months when achieved): head control (10), sitting without support (16), standing (24), and stable walking without support (36). He started to speak at age 39 months and currently is able to speak long sentences, but with unclear pronunciation. Gesell development scale (Chinese version) assessed at age 6 months, 17 months and 50 months showed moderate-mild-moderate impairment in his fine motor domain, severe-moderate-moderate impairment in his adaptive domain, severe-moderate-mild impairment in gross motor domain, severe-bound-moderate impairment in language domain, and profound-mild-mild impairment in his personal social domain. On follow-up he has made developmental progression. His hearing is clinically normal, though BAEP performed at age 6 months and 49 months demonstrated delayed conduction of binaural listening pathways in the brainstem and increased hearing thresholds. Myopia is still present. Brain MRI performed at age 6 months and 13 months showed diffuse myelin deficit in the cerebral white matter, which had resolved at age 50 months (Figure 2). Results of routine investigations and metabolic testing were normal. PMD was suspected clinically, but *PLP1* dosage was normal as were targeted NGS of 115 leukodystrophy-related genes.

**Table S1. Characteristics of three variants in *TMEM63A***

| *Individual* | Gene | CDS | Protein | MAF | CADD | SI | Po | MT | MC | Co | PR | GERP |
|---|---|---|---|---|---|---|---|---|---|---|---|---|
| *1-2* | *TMEM63A* | c.1699G>A | Gly567Ser | - | *35* | *D* | *D* | *D* | *D* | *D* | *D* | *5.45* |
| *3* | *TMEM63A* | c.1385T>A | Ile462Asn | - | *31* | *D* | *D* | *D* | *D* | *D* | *D* | *5.12* |
| *4* | *TMEM63A* | c.503G>A | Gly168Glu | - | *26.9* | *D* | *D* | *D* | *D* | *D* | *D* | *5.91* |

Note: Reference sequence is NM_001258022.1. - indicates the variant absent in population database including 1000G, ExAC or gnomAD; SI: SIFT, Po: Polyphen-2, MT: MutationTaster, MC:M-Cap, Co: Condel, PR: PROVEAN, GERP : GERP++_RS.
